# A role for heavy chain-modification in protecting hyaluronan from free radical fragmentation during inflammation

**DOI:** 10.64898/2026.01.12.699012

**Authors:** Suruchi Poddar, Dorothea A. Erxleben, Rebecca J. Dodd, Heidi L. Reesink, Dixy E. Green, Paul L. DeAngelis, Anthony J. Day, Adam R. Hall

## Abstract

The glycosaminoglycan hyaluronan (HA) is an essential and ubiquitous component of human tissues and biofluids. The only known covalent modification of HA entails the attachment of heavy chains (HC) from the inter-alpha-inhibitor (IαI) family of proteoglycans, forming stable complexes (HC•HA) that arise during inflammation. In some contexts, HC•HA is thought to contribute to pathology, whereas in others it may form part of a protective pathway. However, its exact roles are not fully understood. Here, we report that HC modifications can protect HA from fragmentation by reactive oxygen species (ROS) produced during the inflammatory cascade. Using solid-state nanopore molecular size analysis, we show that HA is highly resistant to degradation from exogenous ROS *in vitro* when in the context of HC•HA complexes, while the unmodified HA polymer is fragmented rapidly under the same conditions. Experiments performed with admixtures of HA and unbound antioxidant proteins – including HC-bearing components – demonstrate the necessity of covalent HC attachment to the polysaccharide for the protection. Finally, we find that HA with high-HC content from ‘inflammatory’ equine synovial fluid has increased resilience to ROS damage compared to low-HC HA from a healthy joint. Collectively, these results demonstrate that covalent HC modification is an effective biological strategy for preserving HA integrity against ROS fragmentation, including in inflammatory conditions.

---

Inflammation is a complex process comprising an elaborate set of interactions between signaling molecules, extracellular matrix (ECM) components, immune cells, and other factors^1^. While it is generally an acute response to injury and infection and serves to protect tissue and restore homeostasis, a growing list of conditions like autoimmune diseases^2^ and obesity^3^ can lead to chronic inflammation that causes widespread damage. For example, the generation of reactive oxygen species (ROS) is a prominent feature of inflammation^4^. While ROS at low levels function as signaling molecules to regulate processes like cellular growth and adhesion, higher accumulation in tissue can induce oxidative stress on cells and cause irreversible damage to DNA or RNA^5^ and degradation of proteins^6^. Such processes are fundamental to the emergence and progression of conditions^7,8^ that include cardiovascular disease, neurodegenerative disease, and several forms of cancer.

Among the myriad biomolecules associated with inflammation is hyaluronan (HA, **Fig. 1a**), a critical glycosaminoglycan (GAG) constituent of the ECM that serves diverse purposes in healthy tissues including hydration, structural support, and maintaining cell adhesion. In inflammation, HA biosynthetic enzymes are upregulated^9^, leading to an increase in overall HA content. However, the molecular weight (MW) of the HA polymer is also a key parameter for its biological activity^10^. In general, it is thought that high-molecular weight HA (HMW-HA) can serve as a signal of tissue integrity^11^, supporting anti-inflammatory, anti-tumorigenic, and wound-healing processes that can include immune cell recruitment and activation^12^. In contrast, the behaviors of fragmented or low-molecular weight HA (LMW-HA) that result from HMW-HA degradation appear to have an opposite function, most likely due to their competion with HMW-HA for HA-receptors and the resultant loss of protective function^10^.

**Figure 1.**
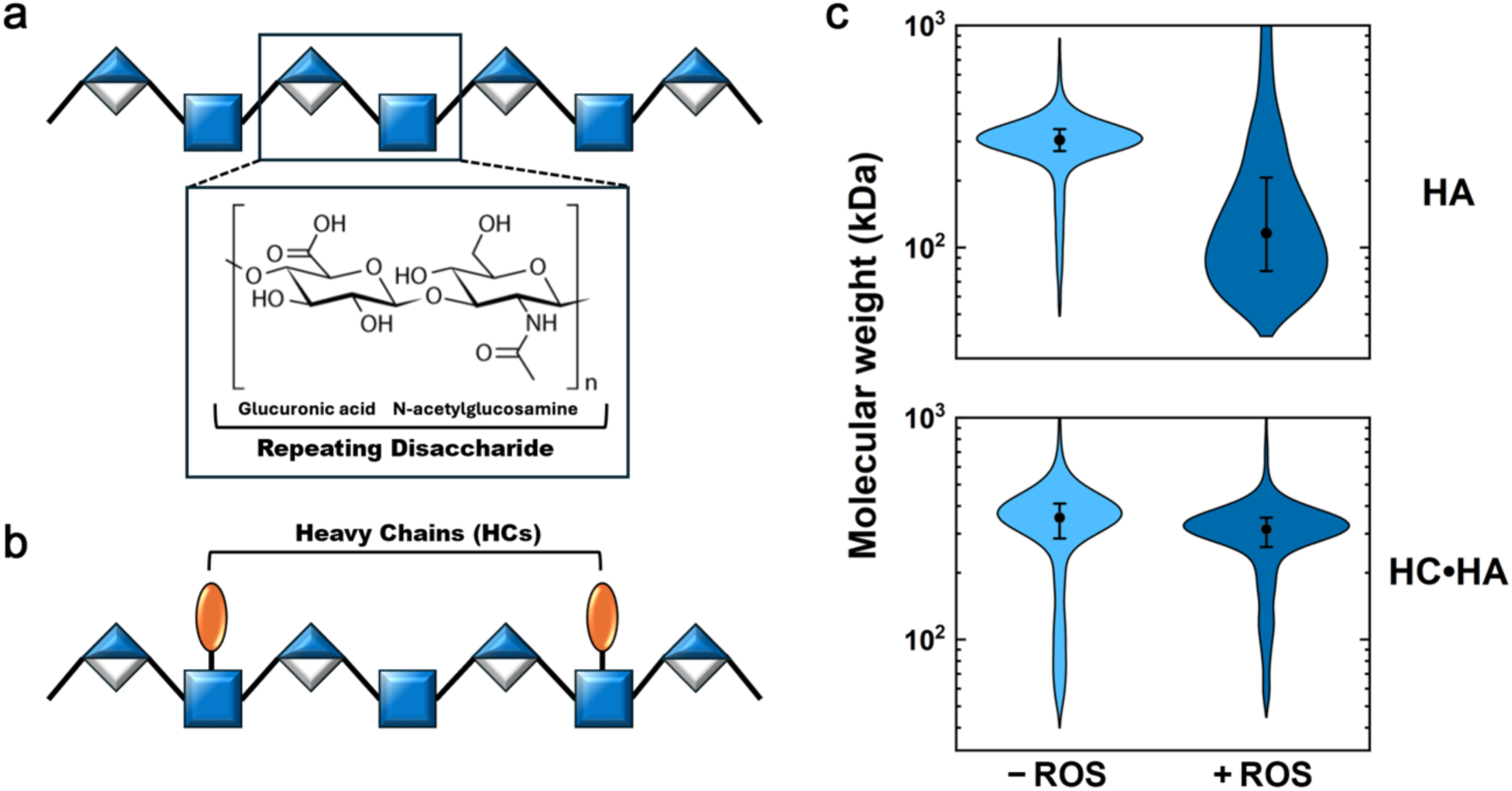
Hyaluronan (HA) and heavy chain-modified hyaluronan (HC•HA) structure and their responses to ROS exposure. (a) Schematic of HA showing the repeating disaccharides of glucuronic acid (semi-filled diamond) and N-acetylglucosamine (filled square). Inset: chemical structure of the HA disaccharide. (b) Schematic of HC•HA complex where an ester bond forms between the C-terminal aspartic acid residue of the HC polypeptide (orange) and a C-6 hydroxyl group of an N-acetyl glucosamine residue. Not to scale. (c) Molecular weight distributions (violin plots) of 321 kDa HA. Upper panel: unmodified HA following in vitro incubation without (-ROS; 305 (+147, −99) kDa) and with (+ROS; 116 (+167, −69) kDa) CuCl_2_/H_2_O_2_. Lower panel: HC-modified HA following in vitro incubation without (-ROS; 355 (+313, −166) kDa) and with (+ROS; 314 (+197, - 121) kDa) CuCl_2_/H_2_O_2_. Listed values indicate geometric median and log-transformed standard deviation.

In addition to the increase in overall HA abundance often observed during inflammation, there is also a concomitant increase in the amount of HA covalently attached via ester bonds^13^ to heavy chain (HC) polypeptides from the inter-α-inhibitor (IαI) family of proteoglycans^10,13–15^. In most cases, IαI is thought to move into tissues from plasma during inflammation, although there is evidence for local biosynthesis in the lung^16^. The esterification of the C-terminal aspartic acid residue of the HC to the C-6 hydroxyl group of an N-acetyl glucosamine residue of HA is driven enzymatically by the inflammation-regulated Tumor Necrosis Factor-Stimulated Gene-6 (TSG-6)^14,15^ protein. HC conjugation is the only known covalent modification to occur on HA, resulting in the formation of so-called ‘heavy-chain hyaluronan complexes’ (HC•HA, **Fig. 1b**).

The association of HA with IαI immunoreactivity was first reported in pathological synovial fluids (SF) in the 1960s^17^ with follow-up studies identifying HC•HA complexes in SF and sera in the context of osteoarthritis and rheumatoid arthritis (RA)^18,19^, in which high levels of TSG-6 are also often present^20^. Since then, HC•HA complexes have been observed across a wide range of inflammatory disorders including asthma^21,22^, inflammatory bowel disease (IBD)^23^, and infection by viruses (*e.g.*, influenza and SARS-CoV-2)^16,24,25^ and nematodes^26^, collectively suggesting their importance in the inflammatory response^10,14,27^. However, the exact roles of HC•HA in these conditions are not clear, such that in some contexts the complexes are implicated in driving disease pathology while in others they likely contribute to tissue protection and repair; for example, it is known that HC•HA promotes leukocyte adhesion^16,28^ that can be either pro- or anti-inflammatory^23,24,29^ and that HCs have the potential to regulate innate immunity via inhibition of the complement system^30,31^. Uncovering additional roles for HC•HA complexes in the inflammatory response continues to be an important area of investigation that could help identify novel therapeutic targets or predictive biomarkers for chronic inflammatory diseases. Here, we demonstrate that covalent HC modifications protect HA from ROS-induced fragmentation *in vitro* using both synthetically-assembled and physiological SF-derived HC•HA complexes. Because such degradation can be detrimental to HA function, it is critical to understand the mechanisms by which it can be avoided and how the beneficial properties of HMW-HA can be maintained.

## HC modification of HA provides protection from oxidative fragmentation

As an unsulfated GAG composed entirely of disaccharides of β-1,4-D-glucuronic acid and β-1,3-N-acetyl-D-glucosamine, HA is highly susceptible to oxidative damage^32^. Indeed, the polysaccharide has long been viewed as a sink for ROS *in vivo* by absorbing the damaging effects of free radicals in place of other important tissue constituents^33,34^. However, such reactions would cause rampant chain scission that could undermine the central roles of HA in tissue structural integrity and function. This situation underscores that extensive damage, especially experienced under chronic ROS exposure, could be detrimental to overall health. Consequently, mechanisms are likely to exist through which the HA polymer itself can be protected to avoid these harmful effects. Due to the association of HC•HA with inflammatory disease and the abundance of ROS in those conditions^7,8,35^, we hypothesized that one role of HC-modification could be to safeguard HA from oxidative fragmentation. In part, this concept was informed by a previous study suggesting that IαI – the source of HCs – in rheumatoid arthritis (RA) SF may be protective against ROS-mediated HA fragmentation^36^. While the IαI was determined to associate with HA, the authors neither suggested nor investigated the formation of a covalent complex. Moreover, two other serum proteins (haptoglobin and α_1_-proteinase inhibitor) were found to be associated with HA in RA SF, but there has been no data to support these proteins attaching covalently to HA. Thus, we first investigated whether HA polymers are resistant to free radical fragmentation *in vitro* when modified covalently with HC.

Solid-state nanopore (SSNP) analysis has proven to be a valuable and ultrasensitive tool for the quantitative single molecule determination of HA size distribution^37,38^, including that of HC-modified HA^39^, and was therefore employed to measure ROS-induced polymer fragmentation. To assess MW changes precisely, unmodified and HC-modified versions of the same quasi-monodisperse HA polymers^40^ were investigated in parallel and compared. To circumvent measurement biases, internal calibration standards were employed for each SSNP (**Supplementary Fig. S1**) and care was taken to strip HCs enzymatically from the HC•HA complexes before analysis^39^. Initially, both specimens yielded a narrow MW distribution centered on their as-produced size of 321 kDa (**Fig. 1c**, light blue; medians: 305 kDa, upper panel, 355 kDa, lower panel). However, following *in vitro* exposure with extrinsic ROS, we observed distinct outcomes. The distribution of the unmodified HA after a 4 h incubation with an established ROS generating system (1 μM CuCl_2_ and 50 mM H_2_O_2_) showed a significant shift towards smaller, more fragmented LMW-HA with a median MW of 116 kDa, or a 64% reduction in size compared to the control (**Fig. 1c**, upper, dark blue). In contrast, the HC•HA treated with identical conditions showed robust protection of the HA from fragmentation (**Fig. 1c**, lower, dark blue), yielding a median MW of 314 kDa, or only a ∼2% reduction compared to control and within experimental error. The mechanism of HA degradation by CuCl_2_ and H_2_O_2_ is through hydroxyl radicals formed by decomposition of the hydrogen peroxide but the effect was not limited to this oxidant pair alone (additional data for the alternative ROS source of ascorbic acid and H_2_O_2_ are shown in **Supplementary Fig. S2**), suggesting that the HC-mediated protection of HA from ROS may be a general property of HC conjugation.

## Covalent linkage of HC to HA is essential to preserve hyaluronan integrity

While these results show that HCs have a protective effect on HA, the nature of that protection is less clear. For example, it is possible that any proximal protein with free radical scavenging activity could provide protection from oxidative fragmentation. In addition, the IαI family of proteoglycans – composed of the core protein bikunin with a single chondroitin sulfate chain to which various HCs are linked – are themselves associated with inflammation. Thus, their presence could provide HCs in the tissue sufficient to consume oxidants without the need for covalent conjugation to HA. To determine the importance of direct HC attachment to the protective process, we next investigated *in vitro* ROS-mediated fragmentation of HA in the presence of ‘bystander’ proteins.

We first assessed the impact of an independent free radical scavenging protein by testing bovine serum albumin (BSA), which has reported antioxidant properties^41^. An admixture of BSA and unmodified HA was produced and subjected to the same CuCl_2_/H_2_O_2_ treatment described above. SSNP size analysis (**Fig. 2**, +BSA) showed a shift in MW from a median value of 287 kDa for the control down to 118 kDa, or a 59% reduction in size. This demonstrated that there was a similar level of oxidative fragmentation to that observed in the absence of BSA and refuted the potential mechanism of an additive antioxidant protein reducing the effective ROS concentration locally to protect the HA. However, features specific to the HC polypeptides could make them particularly efficient at reacting with free radicals in solution. To investigate this possibility, we next used an admixture of unmodified HA and IαI, which typically features two HCs (HC1 and HC2 isoforms) in its structure. After CuCl_2_/H_2_O_2_ exposure, we again observed a substantial shift in the HA MW distribution (**Fig. 2, +**IαI) to a median value of 112 kDa, or a 61% reduction from the control. Therefore, the mere presence of HCs had no observable effect in inhibiting the degradation of the HA.

**Figure 2.**
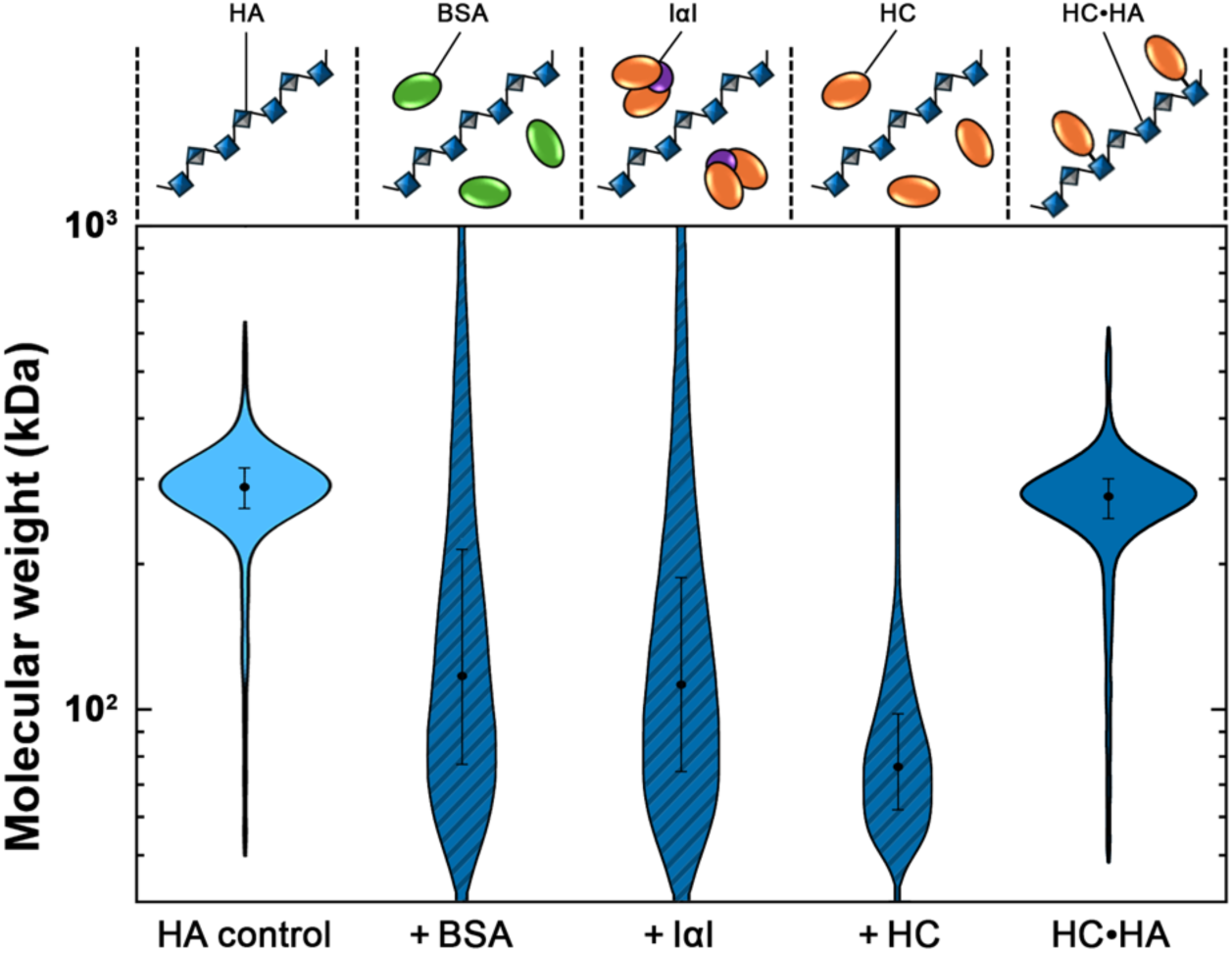
HA fragmentation by ROS in the presence of bystander proteins in comparison to covalent HC•HA. Schematic representations (top) and MW distributions (violin plots) of HA both before (light blue, 287 (+109,-79) kDa) and after in vitro incubation with extrinsic CuCl_2_/H_2_O_2_ (dark blue shaded) in the presence of BSA (118 (+161, −68) kDa), IαI (112 (+143, −63) kDa), or free HCs (76 (+39, −26) kDa), all <u>without</u> covalent attachment to the HA. Listed values indicate geometric median and log-transformed standard deviation. MW distribution of covalent HC•HA after identical ROS incubation is shown (far right, dark blue) for comparison.

Given that HC1, HC2, and bikunin form a globular complex in the context of IαI, the extensive protein-protein interactions between the two HCs^31^ could impact their behavior. Therefore, as a final potential qualifier to our experiments, it is also possible that the HCs within the IαI structure have diminished protective properties relative to their free form. To address this scenario, we repeated the above experiment using IαI after a chemical pre-treatment to release the HCs from their bikunin backbone^15^ prior to ROS exposure (**Supplementary Fig. S3a**). In this way, identical HC content was present as above but in a free state, emulating the HC•HA ensemble without the covalent attachment. Here again, we observed a major change in the HA MW distribution following CuCl_2_/H_2_O_2_ incubation (**Fig. 2**, +HC), yielding a median value of 76 kDa, or a reduction of 74% compared to the control. Note that analysis of the geometric median value does not capture the full change in distribution shape, and so while this shift was larger than that of unmodified HA alone, we do not conclude that free HCs induce additional damage to the HA chain. Similarly, an absence of protection by all bystander proteins was also observed for ascorbic acid and H_2_O_2_ (**Supplementary Fig. S3b**), demonstrating that the fragmentation effect holds beyond this particular oxidative agent. Overall, these results showed conclusively that the covalent attachment of HCs in HC•HA is critical to their HA-protective ability from ROS fragmentation *in vitro*.

What do our experiments reveal about the mechanism of protection by HCs or subsequently about the mechanism of ROS damage in HA? Since HCs are only effective at reducing oxidative fragmentation of HA when they are covalently attached to HA, if the oxidation was a simple local reaction, a free radical generated in solution could first contact either bare HA, where it would sever the glycosidic bond and break the chain, or the associated HC, where it would react instead with the protein and leave the HA polymer intact. However, HC linkages are generally infrequent in native HC•HA complexes; for example, it has been estimated that only 3-5 HCs are attached per ∼2 MDa HA chain in RA synovial fluids^18^. Similarly, the HA in our synthetic HC•HA complexes are also expected to be sparsely decorated with HC proteins, leaving substantial regions of the HA polymer exposed to the environment. This low density would not be expected to protect HA from spontaneous local oxidation and additionally provides no explanation for why covalent HC attachment is critical to the effect. Consequently, we suggest that the degradation of HA by ROS is similar to the oxidation of DNA by radical cations^42^, wherein a radical introduced at a single point in the molecular structure can migrate or ‘hop’ along the molecule until it encounters a potential cleavage site. For DNA, the maximum distance a radical can hop is significant (>20 nm). This distance is dependent on structural and chemical linkage factors and so could conceivably be different for HA. However, free radical hopping without HA bond fision may be less likely if HCs are sufficiently rare, such as in native HC•HA. Therefore, while the robust protection observed in our *in vitro* experiments may have been due to the synthetic HC•HA being built on an HA backbone of only moderate size, it is possible that naturally-occurring, low-level HC-modification on larger sugar chains may not protect HA fully from ROS in tissues and biofluids, where damage could accumulate under chronic exposure.

## Assessing the natural protection of HC•HA in biological fluid

Having established the capability of covalent HC modification to guard HA against oxidative fragmentation, we next investigated the effect on HA in a complex natural biofluid. Here, we compared equine SF specimens collected from animals with advanced (Grade 3) osteoarthritis (OA) to those derived from healthy (Grade 0) animals. There is known to be a large amount of HA per unit volume in SF^43^, where it serves physiologically to hydrate and lubricate the synovium and support smooth joint movement^44,45^. However, HA becomes degraded in OA, losing its lubricating capacity and resulting in increased friction, pain, and cartilage damage^39,46^. Because OA progression has been associated with both HC•HA^23^ and ROS^47^, we identified this as an ideal model system with which to study the protective nature of HC modification in a biological fluid.

We first measured the amounts of HA and HC•HA in each SF specimen using an established ELISA and immunoblotting technique^23^, respectively. These analyses confirmed that, although there was no notable difference in overall HA levels (**Fig. 3a**), HC-modifications were significantly more abundant in the Grade 3 OA SF than in the Grade 0 healthy SF (**Fig. 3b**) and suggested that the former should therefore have the greater ROS-protective capacity based on our observations (*c.f.*, **Fig. 1c**). Each specimen was then divided into two aliquots, one of which served as an untreated control and the other of which was used to test for response to *in vitro* ROS exposure. To emulate physiological ROS buildup, we added CuCl_2_ and H_2_O_2_ directly to the experimental SF specimens up to a final concentration that matched the conditions described above (1 μM and 50 mM, respectively). Following a 4 h incubation, all protein constituents were protease digested to enable unbiased HA MW distributions of each specimen to be determined, including those molecules that had previously featured natural covalent HC modifications. Total HA was then extracted from each sample using a validated affinity-capture approach^48^ and directly assessed with SSNP size analysis.

**Figure 3.**
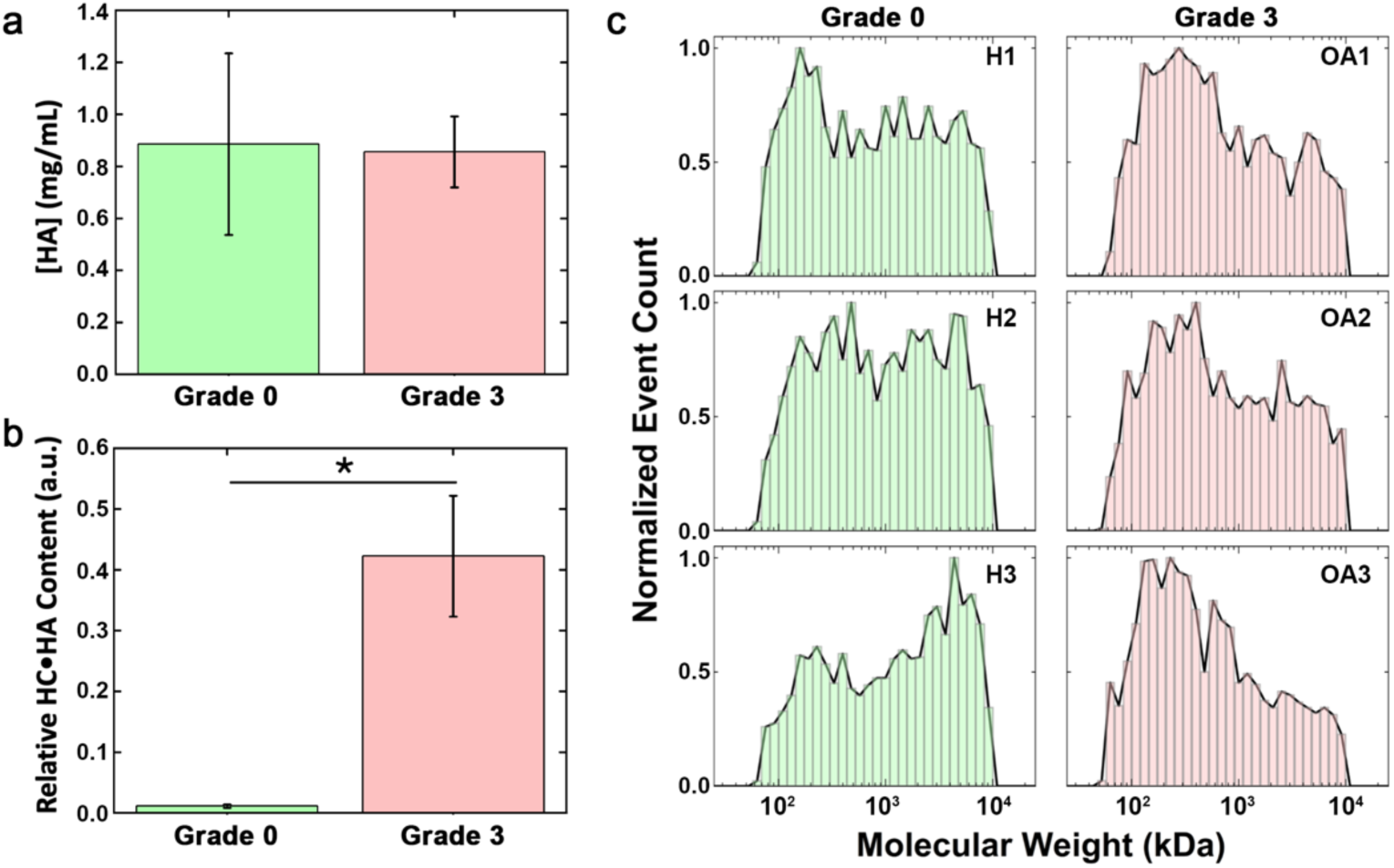
HA and HC•HA concentrations and HA MW distributions in healthy and OA SF. (a) Relative amount of HA and (b) HC•HA from equine SF specimens (Grade 0 healthy, green; Grade 3 OA, red). * indicates p < 0.05 as determined by two-sample t-test. (c) MW histograms of native HA extracted from Grade 0 healthy (H1-H3) or Grade 3 OA (OA1-OA3) equine SF specimens.

Comparing all initial distributions, we observed significant differences between the MW profiles for HA derived from healthy and Grade 3 OA SF (**Fig. 3c**). On average, the healthy HA specimens yielded a median MW value of 989 kDa with a broad log-transformed standard deviation (+2942, −740 kDa), trending more towards HMW-HA than the Grade 3 OA HA with a median of 482 kDa and a narrower log-transformed standard deviation (+1377, −257 kDa); details provided in **Supplemental Table S1**. This finding was in line with previous observations showing that SF HA is degraded during OA^46^ and highlighted that HC modifications do not fully protect native HA from breakdown in chronic exposure. Analyzing the ROS-treated SF, we observed a general reduction in the cumulative HA MW distributions for all samples after CuCl_2_/H_2_O_2_ exposure (**Fig. 4a**), confirming that sufficient undecorated HA remained exposed to enable some fragmentation even in the Grade 3 OA SF. However, the degree of that fragmentation was distinct between cohorts with Grade 0 HA shifting to 371 (+924,-263) kDa and Grade 3 OA HA shifting to 360 (+881,-255) kDa. Considering first the raw change in geometric median MW (**Fig. 4b**), we found that the shift in polymer size was significantly greater for the healthy (low-HC•HA) than for the OA (high-HC•HA) specimens, indicating less protective capacity in the former. Because the baseline HA distributions differed between healthy and Grade 3 OA biospecimens, we also assessed the relative changes in MW as a percentage of their initial values (**Fig. 4c**). Here again, we determined that the fragmentation of the OA HA was significantly less than the healthy HA, corroborating the protective capacity of their comparatively high HC•HA content. As with our measurements on synthetic HC•HA, similar observations were made for ascorbic acid and H_2_O_2_ as a second oxidizing agent (**Supplemental Fig. S4**), suggesting again that fragmentation protection appears to be a role of HC modification against ROS more broadly.

**Figure 4.**
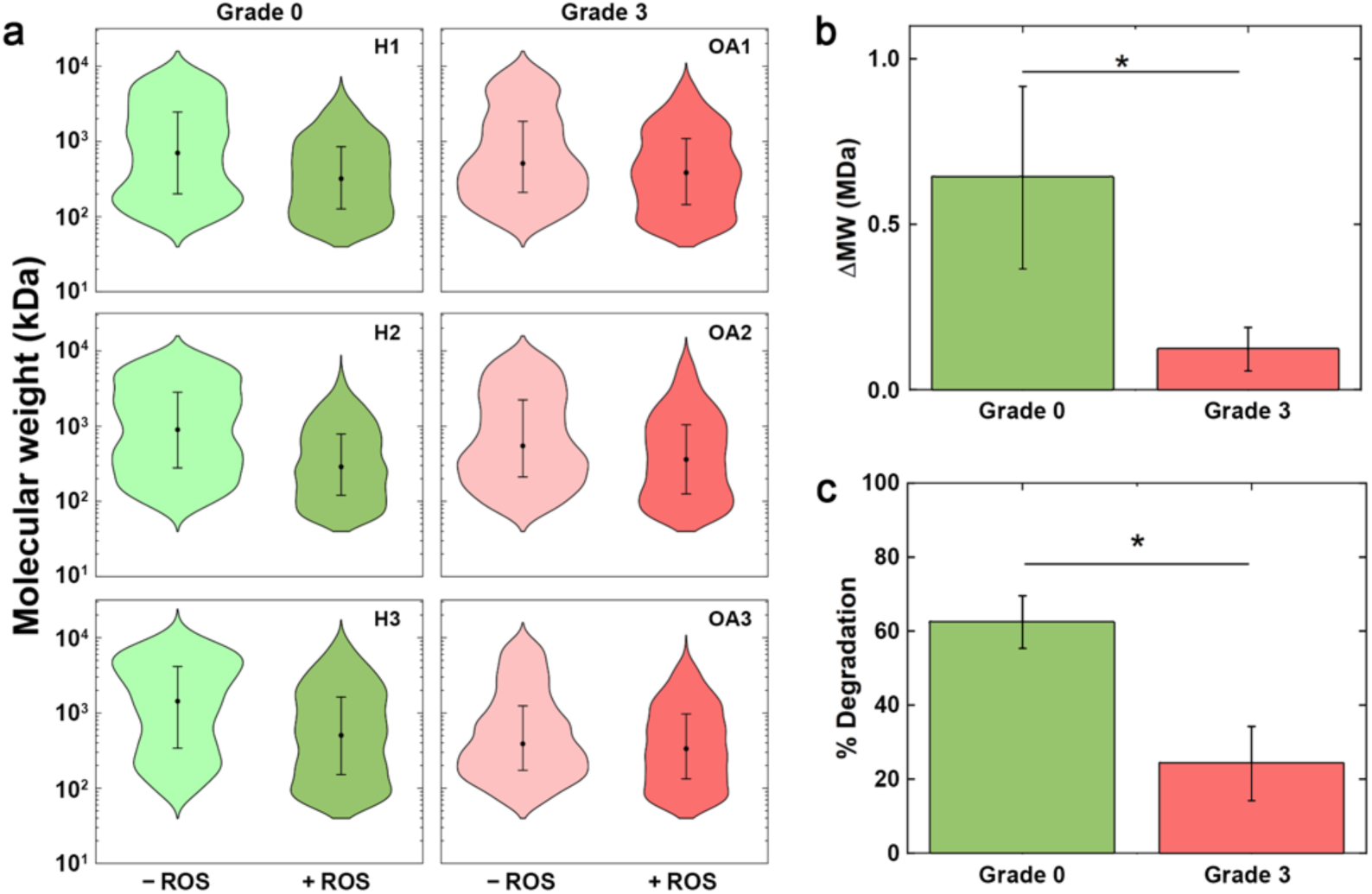
Comparison of ROS-mediated HA fragmentation in healthy and OA synovial fluids. (a) MW distributions (violin plots) for total HA derived from equine SF samples (H1-H3: low-HC content Grade 0; OA1-OA3: high-HC content Grade 3 OA), demonstrating the shift in size of native material (-ROS) after in vitro incubation with CuCl_2_/H_2_O_2_ (+ROS). All individual geometric medians and log-transformed standard deviations are listed in Supplementary Table S2. Comparative analyses of oxidative degradation in SF HA (colors match (a)) after ROS incubation, showing (b) raw geometric median MW change and (c) degradation as a percentage of initial geometric median MW. * indicates p < 0.05 as determined by two-sample t-tests.

## Discussion

HA has also long been viewed as an antioxidant that can provide cells protection from endogenous free radicals through the oxidation of ‘sacrificial’ glycosidic bonds in its chain^33,34^. In some contexts, this breakdown of HA could be an integral mechanism of action; for example, skin HA turns over rapidly with a half life of less than one day^49^, potentially serving to remove and replace HA damaged by environmental radiation and its associated ROS. However, in other important contexts, ROS fragmentation can be detrimental, resulting in a widespread loss of HMW-HA molecules that normally maintain the structural integrity of a healthy ECM and mediate immune regulatory and anti-inflammatory activities. In these settings, oxidative degradation could generate HA fragments that would serve as highly-effective competitive inhibitors of selective HMW-HA-receptor interactions or potentially act as ‘damage’ signals more broadly^10^. Consequently, mechanisms that prevent HA from degrading while still providing tissues with protection from oxidative stress may represent an important physiological function and could be leveraged to limit the deleterious consequences of pathological processes.

We have demonstrated here that HA can be shielded from oxidative fragmentation through the ROS-protective action of HC•HA complexes, wherein the HA is modified covalently with HCs dervived from the IαI family of proteoglycans. Such modifications have previously been shown to promote HA network formation through inter-HC interactions^18,31^ as well as interactions between HCs the octameric protein pentraxin-3 (PTX3)^50,51^. Networks of this kind may serve to increase the effective MW of fragmented HA, partially rescuing its viscoelastic properties in matrices like OA SF where such attributes are critical for joint function. In addition to pathological conditions, the maintenance of viscoelasticity is also important for facilitating normal biological processes, including the expansion of the cumulus-oocyte complex prior to ovulation^52–54^, wherein the essential roles of both ROS^55^ and cumulus matrix integrity suggest that covalent HC protection of HA from oxidative degradation could represent an important mechanism for preserving HMW-HA properties.

Our results further support the mechanistic hypothesis by which ROS interacts with HA networks are analogous to those described previously for other polymers like DNA^56^: free radicals propagate along the polymer chain and can cause damage far from the initial point of contact. We speculate that this process can be mitigated in HC•HA via energy dissipation, leading to oxidative effects within the attached HCs rather than fragmentation of HA. This model may further suggest the existence of a protein-specific quenching mechanism^57^ that could render HC oxidation reversible *in vivo* via the restoration of oxidized HC amino acids through the activity of reductase enzymes. Such action would preserve not only the ROS-protective capacity of HC•HA but also its important crosslink-mediated structural properties and functional activities, such as complement system inhibition^31,53,54^.

One aspect that was not explicitly probed in our study is the minimum HC density needed for efficient chain protection in HC•HA. Given that free radicals have extremely short half-lives (only ∼10⁻⁹ seconds for the hydroxyl (^•^OH) radical^58^), their ranges of diffusion are limited. Therefore, sufficiently sparse HCs would not be expected to prevent oxidative fragmentation effectively. While there is currently no established methodology for determining inter-HC distances explicitly, this may be an important goal for future studies. In general, we expect that optimal HC density in a biological setting must be a balance between providing enough undecorated polymer to maintain the solution properties of HA and enough HCs to prevent rampant degradation. However, the complex 3D nature of HA networks may also play an important role in defining the minimum protective HC density by potentially enabling transmission of radicals between remote regions of the linear molecule brought together by polymer entanglement.

The ROS-protective qualities of HC•HA could provide practical benefits such as maintaining the structure of HA in drug formulations. Indeed, we found that HC•HA MW was preserved in long-term storage relative to naked HA (**Supplemental Fig. S5**). More directly, our observations could also open up new pathways towards improved therapies for inflammation-linked diseases. For example, in diseases such as OA where tissue degradation is progressive, additional (or more active) HC conjugates could provide improved resistance to oxidative damage. Conversely, in some cancers characterized by a dense, HA-rich ECM^59^, the targeting of HC•HA could provide a means by which to both destabilize the tissue matrix and reduce immunosuppressive leukocyte adhesion^28^, potentially improving the efficacy of radio-, chemo-, or immunotherapies.

## Author Contributions

SP performed extractions and nanopore measurements, analyzed the data, contributed to experimental design, and wrote the manuscript. DAE contributed to experimental design. RJD and AJD produced and validated the HC•HA material and contributed to experimental design. DEG and PLD synthesized quasi-monodisperse HA material and contributed to experimental design and analysis. HLR provided the equine SF biospecimens and performed HA and HC•HA quantification. ARH oversaw the project, contributed to experimental design, and wrote the manuscript. All authors contributed to the editing and review of the manuscript.

## Acknowledgements

This work was supported by NIH grants P41EB020594 (ARH) and 5R01GM134226-06 (ARH, PLD, and AJD). We acknowledge funding from Arthritis UK (grant 22277; AJD). RJD was supported by a Wellcome Centre for Cell-Matrix Research Postdoc Grant, funded by the Wellcome Trust (grants 203128/Z/16/Z and 220926/Z/20/Z) and a Wellcome Discovery Award (304200/Z/23/Z to AJD). We also acknowledge the Cleveland Clinic Program of Excellence in Glycoscience (NIH grant P01HL107147) for providing the anti-IαI antibody.

