## Supplemental Methods and Figures for "A role for heavy chain-modification in protecting hyaluronan from free radical fragmentation during inflammation"

### Reagents

Analytical purity $CuCl_2$ dihydrate, 30% aqueous $H_2O_2$, NaOH, HCl, phenol:chloroform:isoamyl alcohol (25:24:1), chloroform, and 100X Tris-EDTA (1 M Tris, 100 mM EDTA, pH 8.0) were all purchased from ThermoFisher Scientific. Ascorbic acid (AA) and lyophilized bovine serum albumin (BSA) were purchased from Sigma Aldrich. *Streptomyces* hyaluronidase was purchased from Millipore Sigma. Inter-alpha-inhibitor (IαI), purified from human serum as described previously[1], was kindly supplied by Jan J. Enghild (Aarhus University, Denmark).

### HA and HC•HA samples

Quasi-monodisperse HA were synthesized chemoenzymatically as described previously[2] (Hyalose, LLC). HA MWs of 111, 545, and 1071 kDa were used to compose solid-state nanopore (SSNP) calibration curves[3] and a 321 kDa sample was employed for subsequent analyses. All MW values were within 5% of the reported mean (polydispersity = 1.001–1.035, as estimated by Size Exclusion Chromatography with Multi Laser Light Scattering). The 321 kDa HA was also employed to produce HC•HA via a transfer reaction[4,5] carried out by incubating 0.94 nM HA with 1.8 μM human serum-derived IαI and 0.28 μM recombinant human TSG-6 (R&D Systems) for 20 hours at 37 °C in a buffer of 20 mM HEPES, pH 7.4, 150 mM NaCl, and 5 mM $MgCl_2$ (100 μL final volume). The reaction was quenched with 5 mM EDTA after the full incubation time. To validate HC transfer, samples from each reaction were analyzed by SDS-PAGE, both with and without pre-digesting HA components with *Streptomyces* hyaluronidase, as described previously[4,5].

**Preparation of oxidative systems and degradation of intact HA**

Two different oxidative systems were used in this study: (a) 1 μM $CuCl_2$ and 50 mM $H_2O_2$ and (b) 20 mM AA and 50 mM $H_2O_2$. After mixing the reactants, HA or HC•HA was immediately introduced and 1X phosphate buffered saline (PBS; 137 mM NaCl, 2.7 mM KCl, 10 mM $Na_2HPO_4$, 1.8 mM $KH_2PO_4$) was used to bring the total volume to 100 μL. Final concentrations of HA (unmodified or as a HC•HA complex) were 37 nM. The mixtures were vortexed briefly and then incubated for 4 h at 37 °C. For experiments investigating bystander effects, BSA and IαI solutions were each prepared by mixing as-received protein in deionized water. Separately, IαI was incubated with 0.1 M NaOH at room temperature for 10 min to liberate HCs (*see* **Figure S3a**), after which 0.1 M HCl was added to neutralize the mixture. Each component was added to an independent aliquot of HA, achieving a final protein concentration of 74 nM – the same 2X excess over HA that was used in the production of HC•HA above – prior to incubation with reactive oxygen species. Following oxidant exposure, HA and HC•HA samples were subjected to ultrafiltration (30 kDa cut-off Amicon Ultra, Millipore Sigma) against deionized water to remove reactants. HC•HA complexes were further treated with a broad-spectrum protease (Proteinase K, Invitrogen) following the manufacturer's instructions to remove HCs from the HA structure and facilitate SSNP size determination. These samples as well as specimens containing bystander proteins were each mixed with an equal volume of phenol:chloroform:isoamyl alcohol, followed by thorough mixing and introduction to a homemade phase-lock tube (2 mL centrifuge tube containing approximately 500 mg of high vacuum grease (Dow Corning)). Samples were then centrifuged for 15 min at 14,000 ×*g* and 20 ˚C to partition the material into an aqueous phase containing HA content and an organic phase containing proteins (and byproducts of proteolysis, where applicable). The same procedure was repeated twice with pure chloroform to remove residual phenol and then subjected to an additional 30 kDa ultrafiltration to remove contaminants prior to analysis. Finally, 10 M LiCl and 100X Tris-EDTA were used to bring each sample to SSNP measurement buffer conditions (6 M LiCl, 1X Tris-EDTA) for size analysis.

**Equine osteoarthritic synovial fluid collection**

Synovial fluid (SF) samples were obtained as described previously[6] using protocols approved by Cornell University's Institutional Animal Care and Use Committee (IACUC protocol numbers 2005-0151 and 2011-0027). Briefly, this study included horses from the Cornell University Equine Hospital undergoing arthroscopic evaluation and treatment of carpal osteochondral fragmentation and/or osteoarthritis (OA) or horses donated for other research projects. No horses had any known history of carpal joint sepsis or intra-articular medication. Joints were scored on the basis of severity of carpal OA as healthy (Grade 0), mild OA (Grade 1), moderate OA (Grade 2), or severe OA (Grade 3) using a previously described scale that includes radiographic assessment with or without additional corroboration from gross or arthroscopic findings[7,8]. Of these, SF was collected from three Grade 0 healthy joints and three Grade 3 OA joints for this study. HA was quantified in each specimen using a commercial enzyme-linked immunosorbent assay (ELISA)-like competitive assay (Echelon Biosciences, K-1200). HC•HA content for each specimen was assessed via an immunoblotting technique described previously[6]. Briefly, in two separate aliquots, 2 µL of SF was diluted in 1X PBS to a final volume of 20 µL. To this was added either 4 µL (0.2 U/µL) of *Streptomyces* hyaluronidase (Millipore Sigma) in 1X PBS or 4 µL of 1X PBS and the mixtures were both incubated at 37°C for 2 h. Each aliquot was analyzed by a Western blot probed with a rabbit polyclonal antibody (A0301) against human IαI (Dako North America, kindly supplied by Vince Hascall, Cleveland Clinic) and a secondary donkey anti-rabbit IgG-HRP (GE Healthcare). Since HC•HA cannot migrate efficiently into gel, the PBS-treated aliquot was used to detect all HCs in the specimen that were not bound to HA, including free HCs and both IαI and Pre-α-inhibitor (PαI) proteoglycans, each of which was represented by a distinct band. The hyaluronidase-treated aliquot assessed all of these HC-containing structures as well as the additional free HCs released from the degraded HC•HA. For each immunoblot, ImageJ software[9] was used to perform densitometric analyses to quantify the free HC band intensity and normalize it with the PαI band, since the latter should remain constant with or without hyaluronidase treatment. The relative amount of HC•HA (a.u.) in each specimen was assessed by subtracting the PBS-treated HC:PαI ratio (background HCs) from the hyaluronidase-digested HC:PαI ratio (background HCs plus formerly HA-bound HCs).

**Affinity extraction of HA from synovial fluid**

HA was extracted following general protocols reported previously[10]. Briefly, equine SF samples (10 μL) were buffer exchanged with 1X PBS using 30 kDa cut-off ultrafiltration. Broad-spectrum protease treatment and solvent extractions/washes were then carried out as described above to partition samples into an aqueous phase (containing HA, other polysaccharides, and nucleic acids) and an organic phase (containing proteins and other byproducts of proteolysis). After decanting the aqueous fraction, HA was then captured and extracted by biomagnetic precipitation using recombinant versican G1 (VG1) domain[11] to bind HA on superparamagnetic beads. For this, streptavidin-conjugated superparamagnetic beads (Dynabeads M-280 Streptavidin, Thermo Fisher Scientific) were first washed according to the manufacturer's directions and then incubated for 1 h at room temperature with biotinylated VG1 (bVG1, Echelon Biosciences) at a ratio of 1 μg bVG1 per 100 μg of beads in 1X PBS. The resulting VG1-beads were washed thoroughly and resuspended into 150 μL aliquots at a concentration of 10 mg/mL in 1X PBS. An aqueous fraction containing HA was added directly to an aliquot of beads and incubated at room temperature for at least 1 h. HA-bound beads were isolated with a magnet and washed to remove any non-specifically bound material. Beads were then incubated with 50 μL of SSNP measurement buffer for 1h, wherein the high ionic strength disrupted the HA-VG1 interaction to elute the HA[12]. The supernatant was retrieved under magnetic partitioning of the beads and used for direct SSNP analysis.

**SSNP measurement and analysis**

Prefabricated SSNPs (P/N: NXPR4002X-16nm-AO-HR and NXPR4001Y-10nm-ABX1) consisting of a single pore in a 20 or 30 nm thick, low-stress silicon nitride membrane were obtained from Norcada, Inc. All pores displayed a linear current-voltage curve with a resistance that yielded a diameter in the range of 6.0−12.0 nm as calculated from an established model[13] that assumes an effective pore thickness equal to 1/3 full membrane thickness due to pore shape. Each device was rinsed with ethanol and water, dried with filtered air, and then treated with air plasma (30 W, Harrick Plasma) for at least 2 min per side before being mounted into a custom 3D-printed (Carbon, Inc.) flow cell. SSNP measurement buffer was introduced to the flow cell to contact each side of the nanopore membrane. Ag/AgCl electrodes were connected to a patch-clamp amplifier (Axopatch 200B, Molecular Devices) and used to both apply voltage and measure current through the nanopore. For HA measurements, isolated HA samples suspended in 10 μL of SSNP measurement buffer were loaded on one side of the pore. A 300 mV bias was then applied, and the transmembrane current was recorded at a rate of 200 kHz using a 100 kHz four-pole Bessel filter. Data were collected and analyzed with a custom LabVIEW program (National Instruments). An additional 5 kHz low-pass filter was applied during analysis. Molecular translocations were marked by temporary reductions in the ionic current and these 'events' were identified using a threshold of 5 standard deviations of the root-mean-square noise. The event charge deficit (ECD) of each event (*i.e.*, the integrated event area, dependent on both translocation duration and amplitude) was correlated to its corresponding MW[14] using an internal calibration curve (**Supplementary Fig. S1**) produced by measuring a mixture of three quasi-monodisperse HAs (111, 545, and 1071 kDa, as described above) on each SSNP prior to HA sample analysis. Gaussian centers of the three peaks yielded a linear relationship with ECD for calibration. Events with ECD values corresponding to MWs between 50 kDa and 10 MDa were considered for further analyses.

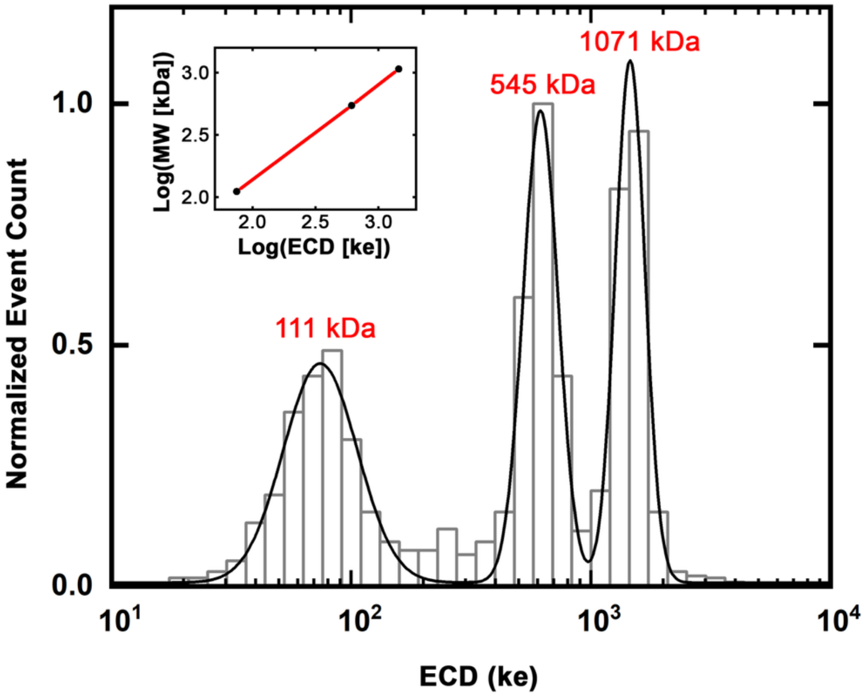


**Supplementary Figure S1.** Example SSNP ECD histogram showing the discernable peaks of a mini-ladder composed of three independent, quasi-monodisperse HAs (111, 545, and 1071 kDa, respectively). The peak centers from the multipeak Gaussian fit (black line) are used to define an internal calibration curve (inset) with which SSNP signals from subsequent HA measurements can be converted to MW on a molecule-by-molecule basis. The red line on the inset is a linear fit on a log-log scale as described in Ref. 14

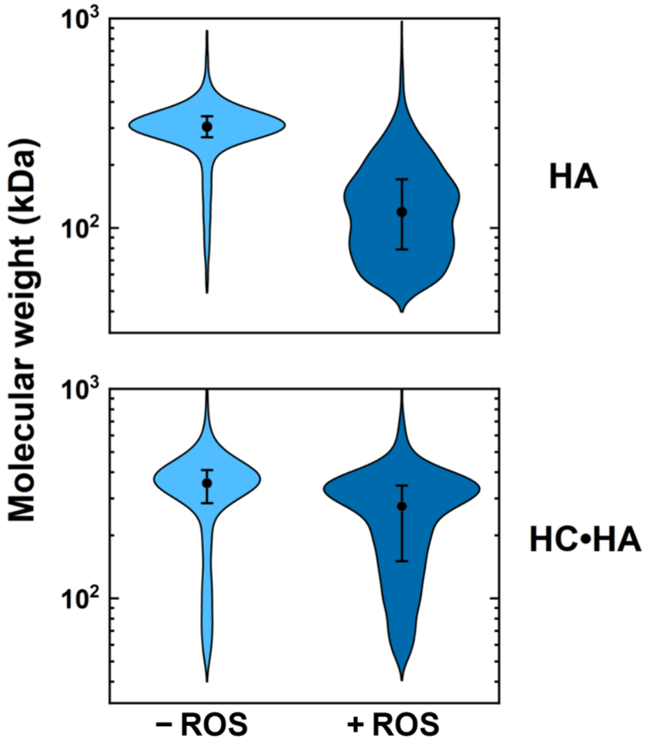


**Supplementary Figure S2.** Molecular weight distributions (violin plots) of 321 kDa HA. Upper panel: unmodified HA following *in vitro* incubation without (-ROS; 305 (+147, -99) kDa) and with (+ROS; 119 (+83, -49) kDa) AA/$H_2O_2$. Lower panel: HC-modified HA following *in vitro* incubation without (-ROS; 355 (+313, -166) kDa) and with (+ROS; 275 (+234, -126) kDa) AA/$H_2O_2$. Listed values indicate geometric median and log-transformed standard deviation.

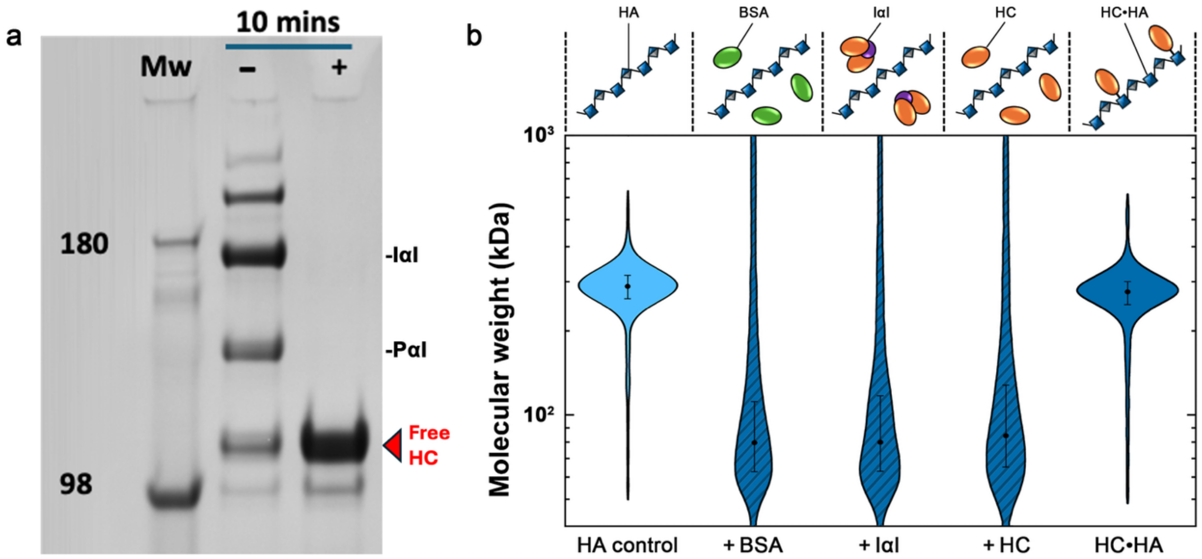


**Supplementary Figure S3.** (a) SDS-PAGE of IαI before (-) and after (+) a 10 min treatment with NaOH, demonstrating the liberation of HCs by hydrolysis of the ester bonds linking HCs to the chondroitin sulfate chain of bikunin in IαI. Initial material contains pre-α inhibitor (PαI). Upper bands are likely IαI with 3 and 4 HCs attached, respectively. (b) Schematic representations (top) and MW distributions (violin plots) of HA both before (light blue, 287 (+109,-79) kDa) and after *in vitro* incubation with extrinsic AA/$H_2O_2$ (dark blue shaded) in the presence of BSA (79 (+56, -33) kDa), IαI (79 (+59, -34) kDa), or free HCs (84 (+80, -41) kDa), all without covalent attachment to the HA. Listed values indicate geometric median and log-transformed standard deviation. MW distribution of covalent HC•HA (dark blue; 275 (+106, -77) kDa) after identical ROS incubation is shown (right, dark blue) for comparison.

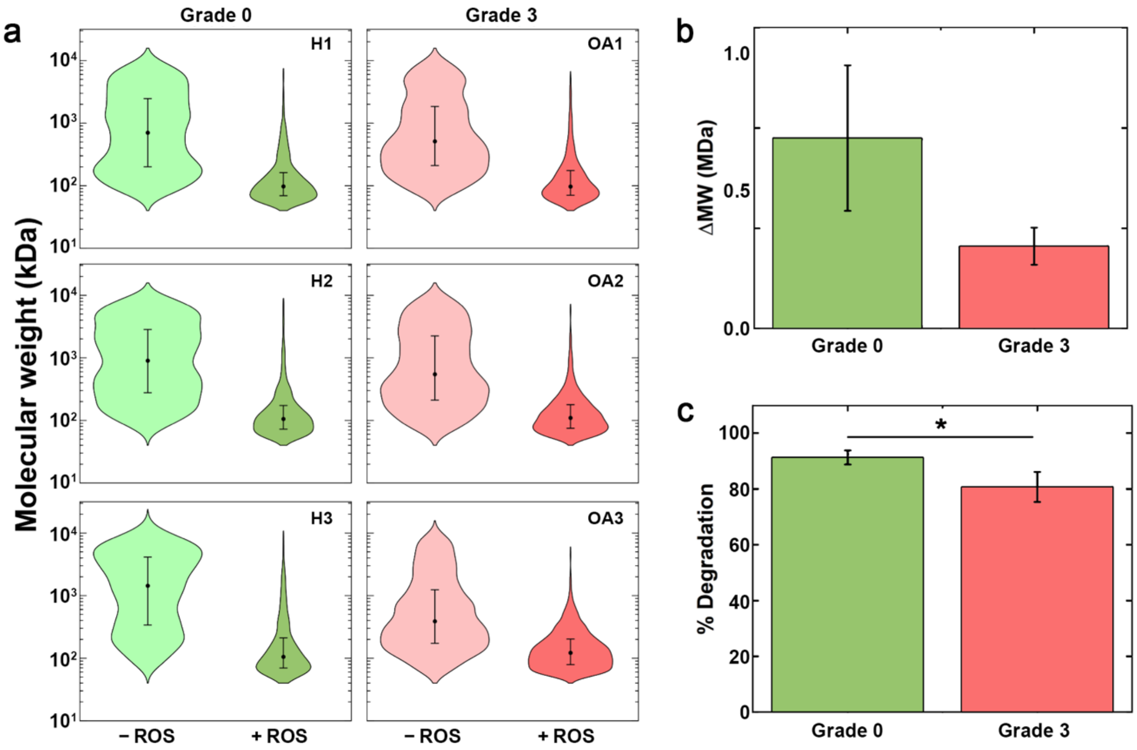


**Supplementary Figure S4.** (a) MW distributions (violin plots) for total HA derived from equine SF samples (H1-H3: low-HC content, Grade 0; OA1-OA3: high-HC content, Grade 3 OA) demonstrating the shift in the distributions of native material (-ROS, light) after *in vitro* incubation with AA/$H_2O_2$ (+ROS, dark). All individual geometric medians and log-transformed standard deviations are listed in Supplementary Table S3. (b,c) Comparative analyses of oxidative degradation in SF HA (colors match (a)) after ROS incubation, showing (b) raw MW change and (c) degradation as a percentage of initial MW. * indicates $p < 0.05$ as determined by two-sample t-test.

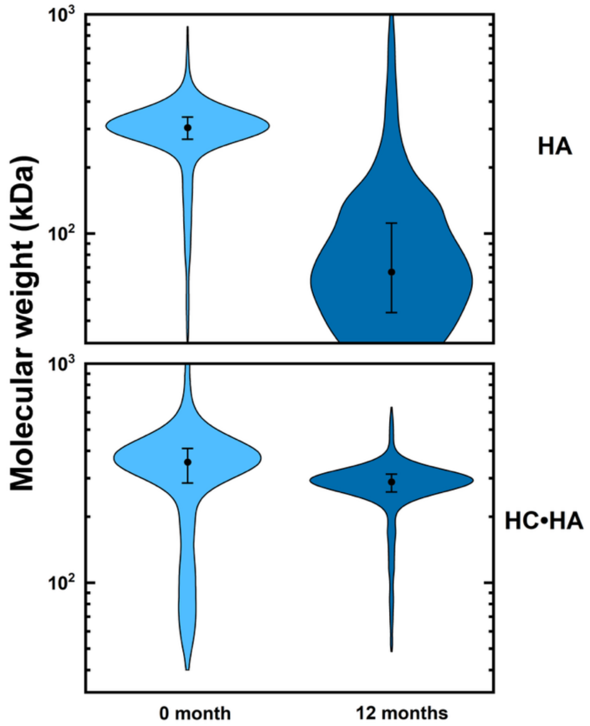


**Supplementary Figure S5.** MW distributions of HA (top) and HC•HA (bottom) measured before and after 12 months of storage at 4 $^{o}$C. Undecorated HA is heavily degraded whereas the HC•HA maintains its structure, indicating that ROS fragmentation is a major mechanism of HA degradation during storage and can be reduced through HC modification.

|  | Sample | # Events | Geometric Mean | Geometric Median | Standard Deviation |
|---|---|---|---|---|---|
| *Units* |  |  | *kDa* | *kDa* | *kDa* |
| **Grade 0** | **H1** | 1748 | 727 | 702 | +2169 |
|  |  |  |  |  | -530 |
|  | **H2** | 1989 | 884 | 899 | +2567 |
|  |  |  |  |  | -666 |
|  | **H3** | 1960 | 1122 | 1365 | +4090 |
|  |  |  |  |  | -1023 |
| **Grade 3** | **OA1** | 1829 | 637 | 511 | +1458 |
|  |  |  |  |  | -378 |
|  | **OA2** | 1954 | 668 | 546 | +1619 |
|  |  |  |  |  | -408 |
|  | **OA3** | 2065 | 501 | 389 | +1056 |
|  |  |  |  |  | -284 |

**Supplementary Table S1.** Descriptive statistics for native (untreated) HA distributions from Grade 0 healthy and Grade 3 OA equine SF specimens used in this study**.**

| | Sample | # Events | Geometric Mean | Geometric Median | Standard Deviation |
|---|---|---|---|---|---|
| *Units* | | | *kDa* | *kDa* | *kDa* |
| **Grade 0** | **H1** | 1408 | 336 | 320 | +667 |
| | | | | | -216 |
| | **H2** | 1405 | 313 | 288 | +600 |
| | | | | | -194 |
| | **H3** | 1539 | 508 | 504 | +1506 |
| | | | | | -378 |
| **Grade 3** | **OA1** | 1545 | 401 | 384 | +920 |
| | | | | | -271 |
| | **OA2** | 1414 | 388 | 360 | +946 |
| | | | | | -261 |
| | **OA3** | 1333 | 366 | 334 | +779 |
| | | | | | -234 |

**Supplementary Table S2.** Descriptive statistics for HA distributions from Grade 0 healthy and Grade 3 OA equine SF specimens used in this study following *in vitro* $CuCl_2/H_2O_2$ incubation**.**

| | Sample | # Events | Geometric Mean | Geometric Median | Standard Deviation |
|---|---|---|---|---|---|
| *Units* | | | *kDa* | *kDa* | *kDa* |
| **Grade 0** | **H1** | 1080 | 122 | 97 | +121 |
| | | | | | -54 |
| | **H2** | 1099 | 129 | 105 | +138 |
| | | | | | -60 |
| | **H3** | 1008 | 145 | 105 | +181 |
| | | | | | -66 |
| **Grade 3** | **OA1** | 966 | 128 | 97 | +130 |
| | | | | | -56 |
| | **OA2** | 1356 | 128 | 110 | +122 |
| | | | | | -58 |
| | **OA3** | 1600 | 137 | 122 | +128 |
| | | | | | -62 |

**Supplementary Table S3.** Descriptive statistics for untreated HA distributions from Grade 0 healthy and Grade 3 OA equine SF specimens used in this study following *in vitro* AA/$H_2O_2$ incubation**.**
